# A Multimodal Atlas of Tumor Metabolism Reveals the Architecture of Gene-Metabolite Co-regulation

**DOI:** 10.1101/2022.11.23.517549

**Authors:** Elisa Benedetti, Eric Minwei Liu, Cerise Tang, Fengshen Kuo, Mustafa Buyukozkan, Tricia Park, Jinsung Park, Fabian Correa, A Ari Hakimi, Andrew Intlekofer, Jan Krumsiek, Ed Reznik

**Author notes:** Correspondence to: Jan Krumsiek, Ed Reznik. Equal contribution (Elisa Benedetti, Eric Minwei Liu, Cerise Tang).

## Abstract

Tumor metabolism is controlled by coordinated changes in metabolite abundance and gene expression, but simultaneous quantification of metabolites and transcripts in primary tissue is rare. To overcome this limitation and study gene-metabolite coregulation in cancer, we assembled the Cancer Atlas of Metabolic Profiles (cAMP) of metabolomic and transcriptomic data from 988 tumor/normal specimens spanning 11 cancer types. Meta-analysis of the cAMP revealed two classes of Gene-Metabolite Interactions (GMIs) that transcended cancer types. The first corresponded to a small number of gene-metabolite pairs engaged in direct enzyme-substrate interactions, identifying putative metabolite-pool-size-controlling genes. A second class of GMIs represented a small number of hub metabolites, including quinolinate and NAD+, which correlated to many genes specifically expressed on immune cell populations. These results provide evidence that gene-metabolite coregulation in human tissue arises, in part, from both mechanistic interactions between genes/metabolites, and from metabolic remodeling in specific immune microenvironments.

## Introduction

Coordinated changes of genetically encoded metabolic enzymes and transporters, and the metabolites they act on, underpin diverse cancer-associated phenomena, including tumorigenesis^1^, pluripotency^2,3^, the onset of drug resistance^4–6^, and the modulation of immune responses^7–11^. However, in spite of the high value of joint profiling of metabolites and gene expression/protein levels, previous large-scale studies of tumor metabolism have overwhelmingly focused on the analysis of gene expression data^12^. Conversely, the few instances of multimodal metabolomic and transcriptomic profiling of human tumor specimens have largely been performed in disparate, unrelated studies by a multitude of research teams^13–22^. Integration of metabolomic datasets produced in different patient cohorts is challenging due to technical batch effects and the semi-quantitative nature of untargeted metabolomic data (reported in arbitrary units of relative abundance). Thus, both the scarcity of multimodal metabolomic/transcriptomic data from tissue specimens and the challenges of harmonizing available datasets fundamentally impede the discovery of recurrent, coordinated changes in metabolic gene expression and metabolite abundance across cancers.

Tumors from diverse cancer types differ in their cell-type composition, vascularization, and other factors ultimately influencing metabolism. Yet, they share a convergent set of metabolic alterations^23–27^. For example, several meta-analyses of the tumor metabolic transcriptome have identified recurrent up-regulation of genes in one-carbon metabolism and oxidative phosphorylation across cancer types^26,28,29^. Analogously, meta-analyses of metabolomics data have demonstrated that numerous central carbon metabolites (*e*.*g*., lactate) and effector metabolites (*e*.*g*., kynurenine) are at higher abundance in tumor tissue compared to normal tissue across many cancer types^23,30^. These studies have illustrated the power of meta-analysis for distilling highly recurrent metabolic phenotypes from heterogeneous data but have left unresolved the question of how transcription of metabolic genes and metabolite abundance are coordinated and ultimately shape tumor physiology.

To systematically investigate gene-metabolite coregulation in cancer, we assembled, harmonized, and integratively analyzed metabolomics and transcriptomics profiles from 988 primary tumor and matched adjacent normal tissue collected in 15 independent studies covering 11 cancer types. The preprocessed and harmonized data constitute the Cancer Atlas of Metabolic Profiles (cAMP), representing what is to our knowledge the largest harmonized dataset of multimodal metabolomic and transcriptomic data on primary tumor specimens. The cAMP is publicly available for download and can be interactively explored at https://rezniklab.shinyapps.io/cAMP-shiny-app/. Leveraging the diversity of diseases in our dataset, we designed a concordance-based statistical meta-analysis approach to discover instances of Gene-Metabolite Interactions (GMIs) that transcended cancer type. This revealed two distinct classes of GMIs: First, we identified a small number of strong interactions between enzymes and metabolites involved in the same or subsequent reactions (“proximal” GMIs), suggesting that these enzymes are the primary determinants of their respective metabolite pool sizes. A second group of GMIs consisted of a small number of metabolites broadly correlated to large numbers of genes. Interestingly, this second class of GMIs were enriched for genes specifically expressed in immune cells, and for metabolites related to nicotinamide adenine dinucleotide (NAD+), a pleiotropic metabolite which acts both as a central cofactor in metabolism^31^ and as a signaling molecule influencing cell identity^32^. Taken together, these findings suggest that gene-metabolite co-regulation in tumors is controlled, in part, by two complementary phenomena: the expression of enzymes with strong control over metabolite pool size, and the presence of specific cell populations in the tumor microenvironment with characteristic metabolomic profiles.

## Results

### A Cancer Atlas of Metabolic Profiles (cAMP) reveals the architecture of gene-metabolite coregulation across cancers

Since metabolomic profiling has so far been excluded from large multi-modal tumor profiling projects *(e*.*g*., in the TCGA^33^), there is no unified resource of metabolomic/transcriptomic data in the cancer research field. However, several groups have independently produced and released matched metabolomic/transcriptomic data in diverse cancer types^13–22^. We combined these datasets with several in-house studies to create a comprehensive collection of 988 samples (764 tumor sample and 224 adjacent normal samples) across 11 different cancer types, covering 15 datasets, which we called the Cancer Atlas of Metabolic Profiles (cAMP) (**Table 1** and **Figure 1A**). The overall collection includes a total of more than 40,000 unique transcripts and almost 2,500 unique metabolites. To maximize comparability across these heterogeneous studies, we applied a unified workflow to process RNA expression from microarray and RNA sequencing data, harmonize metabolite names and annotations, and standardize data normalization and preprocessing (see **Methods**). The cAMP represents an unprecedented resource to investigate the coregulation of metabolite levels and gene expression at scale across diverse lineages of human cancers and normal tissues.

**Table 1.** Dataset Overview.

| **Cohort** | **Cancer Type** | **Reference** | **Samples**<br>**(T / N)** |
| --- | --- | --- | --- |
| BRCA1 | Breast Cancer | Terunuma et al. 2014<sup>19</sup> | 61 / 47 |
| BRCA2 | Breast Cancer | Tang et al. 2014<sup>20</sup> | 18 / -- |
| COAD | Colon Adenocarcinoma | Satoh et al. 2017<sup>18</sup> | 37 / 39 |
| DLBCL | Diffuse Large B-cell Lymphoma | Calvo-Vidal, Zamponi et al. 2021<sup>17</sup> | 62 / -- |
| GBM | Glioblastoma | Wang et al. 2021<sup>16</sup> | 74 / 6 |
| HürthleCC | Hürthle Cell Carcinoma | Ganly, Liu et al. 2022<sup>86</sup> | 28 / 3 |
| HCC | Hepatocellular Carcinoma | Chaisaingmongkol et al. 2017<sup>15</sup> | 54 / -- |
| ICC | Intrahepatic Cholangiocarcinoma | Chaisaeng Mongkol et al. 2017<sup>15</sup> | 86 / -- |
| OV | High Grade Serous Ovarian Cancer | Gentric et al. 2019<sup>14</sup> | 45 / -- |
| PDAC | Pancreas Adenocarcinoma | Zhang et al. 2013<sup>21</sup> | 27 / 12 |
| PRAD | Prostate Adenocarcinoma | Penney et al. 2020<sup>13</sup> | 91 / 46 |
| ccRCC1 | Clear Cell Renal Carcinoma | Hakimi et al. 2016<sup>22</sup> | 32 / -- |
| ccRCC2 | Clear Cell Renal Carcinoma | in-house | 30 / -- |
| ccRCC3 | Clear Cell Renal Carcinoma | in-house | 67 / 47 |
| ccRCC4 | Clear Cell Renal Carcinoma | in-house | 52 / 24 |

**Figure 1.**
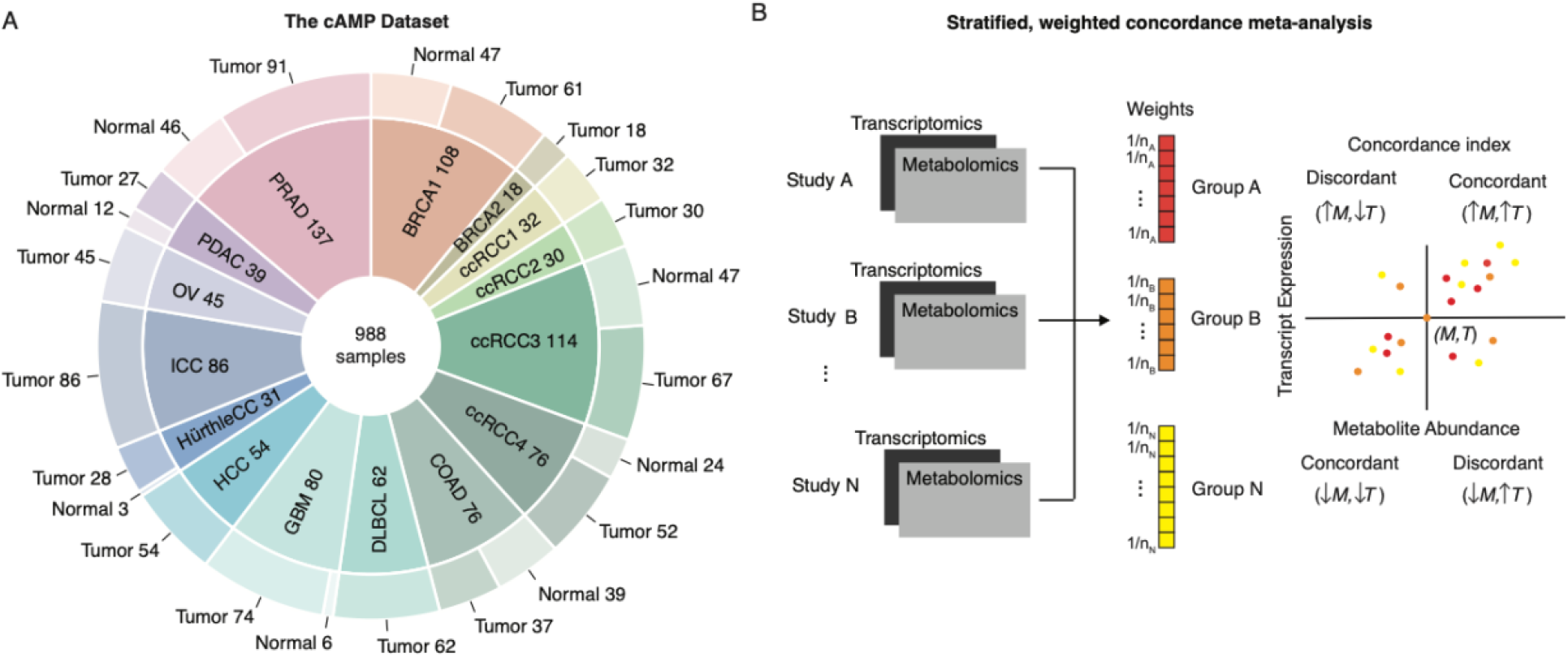
Summary of the Cancer Atlas of Metabolic Profiles (cAMP). (A) The cAMP integrates metabolomic and transcriptomic data from 15 datasets covering 11 different cancer types, covering both tumor and normal tissue. Cancer type abbreviations, BRCA1/BRCA2: breast cancer, COAD: colon adenocarcinoma, DLBCL: diffuse large B-cell lymphoma, GMB: glioblastoma, HürthleCC: Hürthle cell carcinoma, HCC: hepatocellular carcinoma, ICC: intrahepatic cholangiocarcinoma, OV: high-grade serous ovarian cancer, PDAC: pancreas adenocarcinoma, PRAD: prostate adenocarcinoma, ccRCC1/ccRCC2/ccRCC3/ccRCC4: clear cell renal carcinoma. (B) Overview of non-parametric concordance meta-analysis. For a given transcript (“T”) and metabolite (“M”) pair, all measurements from all datasets are considered and weighted according to sample size. The concordance is a non-parametric measure of bivariate association between T and M that can be applied in meta-analysis across multiple datasets.

We interrogated the cAMP dataset for recurrent covariation between genes and metabolites across datasets and cancers. Such gene-metabolite coregulation could emerge via numerous mechanisms including direct metabolic interactions, *e*.*g*., via expression changes of a rate-limiting enzyme, or by the accumulation/depletion of metabolites as part of a broader phenotype, such as a cytotoxic immune response^34–36^. While each cancer type is likely to demonstrate its own unique pattern of transcriptomic and metabolomic changes, the cAMP enables discovery of metabolomic/transcriptomic covariation that transcends diseases. To identify cancer-type-agnostic metabolite/transcript covariation across cAMP datasets in a statistically principled manner, we developed a concordance-based meta-analysis approach (**Figure 1B, Methods**). Concordance is a non-parametric measure of correlation, similar to Kendall’s Tau, which enables the identification of consistently positive or negative gene-metabolite associations across datasets^37^. We focused on the 276 metabolites that were quantified in more than half of the tumor datasets (at least 8 of 15 tumor datasets) and the 16,082 genes that were quantified in all 15 studies. Out of all possible gene-metabolite pairs (276 metabolites x 16,082 genes = 4,438,632 pairs), a total of 22,619 pairs (0.51%) were significantly correlated after multiple testing correction at 0.01 FDR (**Figure 2A**). This included 269 metabolites (∼97%) and 7,987 genes (∼50%) participating in at least one significant association (**Table S1**), which we refer to as Gene-Metabolite Interactions (GMIs).

**Figure 2.**
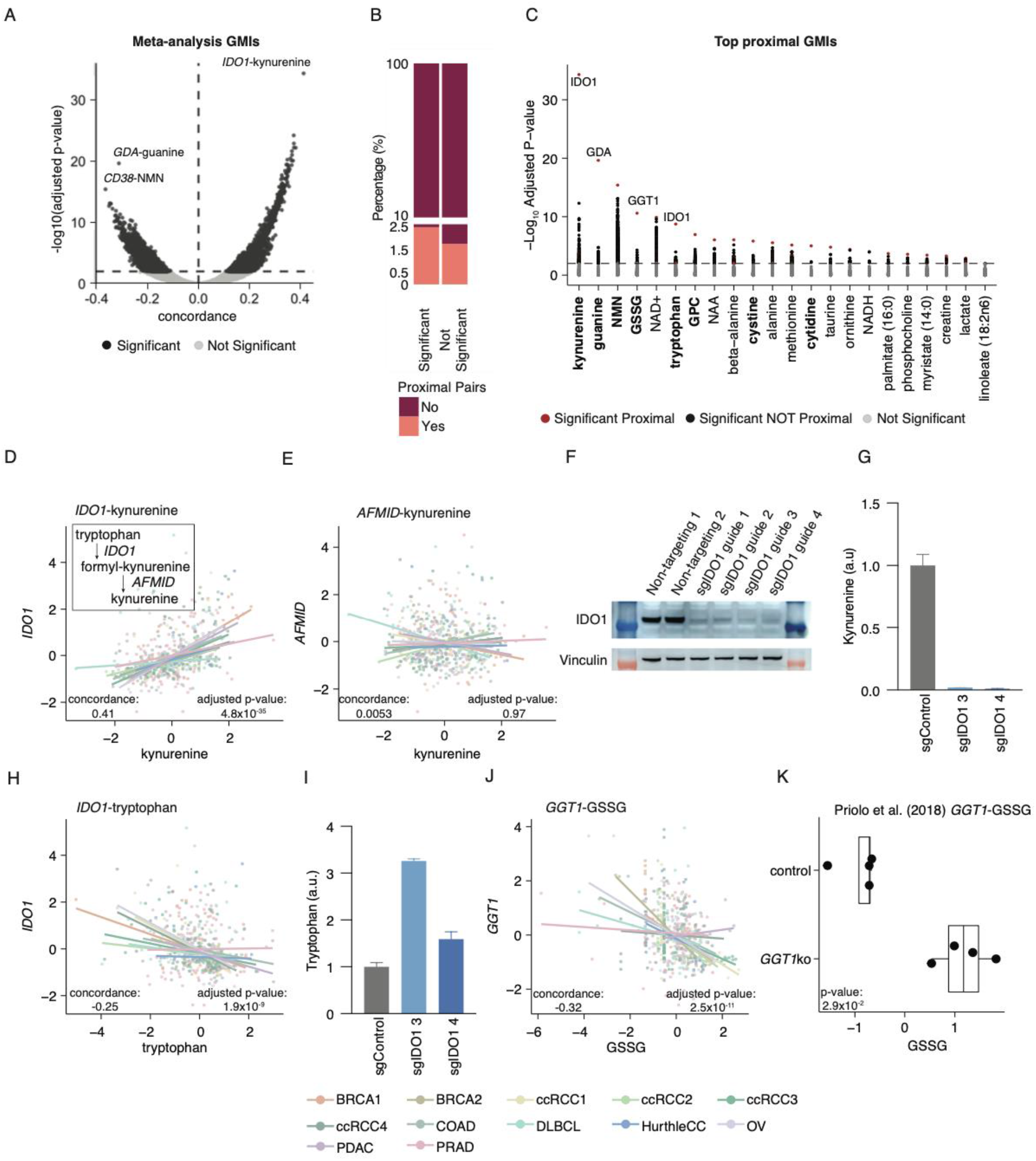
Meta-analysis across the cAMP captures lineage-transcending gene-metabolite interactions. (A) Volcano plot of the gene-metabolite interactions (GMIs) computed between the 16,082 genes present in all datasets, and the 276 metabolites present in at least 8 of our 15 tumor cohorts. The x-axis indicates the scaled concordance value, where values above 0 indicate positive association and values below 0 indicate negative association. The y-axis represents the corresponding -log_10_ FDR-adjusted p-value. The horizontal line indicates the significance cutoff of 0.01 FDR. Light gray dots indicate non-significant gene-metabolite pairs, and black dots indicate significant pairs. Three top hits have been highlighted. (B) Statistically significant GMIs are enriched for proximal interactions, but proximal interactions nevertheless constitute a minority of all statistically significant GMIs. (C) Proximal GMI prioritization. GMIs for the 22 metabolites whose strongest GMI was proximal (distance less or equal 2). For each metabolite, we ranked genes by their statistical significance. Red and black dots indicate proximal and non-proximal genes significantly associated with the corresponding metabolite, respectively, while gray dots indicate genes with non-significant associations. Bold metabolites exhibit a large gap between the dominant GMI and all other GMIs for a metabolite. (D-E) Scatter plots of the association between kynurenine levels and two proximal genes (*IDO1*, Panel D; *AFMID*, Panel E). Metabolite abundances were scaled within each dataset to be displayed together. (F) CRISPR-CAS9 mediated knockdown of *IDO1* depletes *IDO1* protein levels in HCT116 cells. (G) Kynurenine levels are depleted upon knockdown of *IDO1* in HCT116 cells. (H) Scatter plots of the association between tryptophan levels and *IDO1* in the cAMP. (I)Tryptophan levels increase upon *IDO1* knockdown. (J) Scatter plot of oxidized glutathione (GSSG) and GGT1 (K) Validation of the GSSG and GGT1 relationship based on the study from Priolo et al.^42^.

### A small number of strong GMIs correspond to direct enzyme-substrate interactions

Among the statistically significant GMIs (**Figure 2A**), we noted that the strongest positively correlated GMI (*IDO1-*kynurenine, adjusted p-value: 4.82×10^−35^ and the two strongest negatively correlated GMIs, *GDA*-guanine (adjusted p-value: 2.31×10^−20^) and *CD38*-nicotinamide ribonucleotide (NMN) (adjusted p-value: 3.90×10^−16^) corresponded to “proximal” metabolic interactions, which are interactions between an enzyme and its direct or nearly direct substrate/product. For example, *IDO1* catalyzes the catabolism of tryptophan to N-formyl-kynurenine, which is subsequently metabolized to kynurenine^38^, and both *CD38* and *GDA* directly degrade the metabolites guanine and NMN, respectively^39,40^. We confirmed that statistical significance for these 3 GMIs was likely driven by several cancer types rather than a single dataset with very strong associations (**Figure S1**).

The direct biochemical relationship between the three GMIs above raised the possibility that functional proximity between enzymes and their substrates/products might underlie a large fraction of GMIs. To test this, we systematically computed the distance between all gene-metabolite pairs using the highly curated Human-1 metabolic network model^41^ (see **Methods**). Although statistically significant GMIs were enriched for proximal interactions relative to non-significant gene-metabolite pairs (odds ratio: 1.42, Fisher’s exact test p-value: 3.09 x10^−15^), proximal interactions themselves constituted only a small fraction of the total ensemble of statistically significant GMIs (2.5%, 565/22,619) (**Figure 2B**). Thus, while several of the strongest GMIs arose from proximal interactions, gene-metabolite proximity was a weak determinant of the full GMI landscape.

To further investigate the above observations about the strength of specific proximal GMIs and the overall relationship between metabolic proximity and GMIs, we investigated the relative strength of different GMIs affecting a common metabolite. We focused on the 22 metabolites whose strongest GMI was proximal, covering diverse molecules involved in nucleotide metabolism (guanine, cytidine), cofactor metabolism (NAD+), redox metabolism (cystine, GSSG), and other pathways (**Figure 2C**). Interestingly, we found that for 8 out of 22 metabolites (kynurenine, guanine, NMN, GSSG, tryptophan, GPC, cystine, and cytidine), a large gap existed between the most significant GMI and the second-highest correlating transcript. This gap suggested that the pool size of these metabolites was strongly controlled, in a lineage-agnostic manner, by a single, dominant gene. Consequently, we hypothesized that targeted genetic knockdown of dominant GMIs for these 8 metabolites would have a higher likelihood of producing significant changes in pool size than for other metabolites with multiple, comparably strong GMIs, each of which might control the pool size of the metabolite (**Figure 2C**, bold).

We sought to functionally validate a subset of these predicted, metabolite pool-controlling genes. First, we investigated the association between *IDO1* and two metabolites, kynurenine and tryptophan. *IDO1* converts tryptophan to N-formylkynurenine, which is subsequently catabolized to kynurenine by *AFMID*. We observed that kynurenine levels were strongly associated with *IDO1* (**Figure 2D**) but not *AFMID* (which acts directly to produce kynurenine, **Figure 2E**) expression across the cAMP, consistent with previously published results from the Cancer Cell Line Encyclopedia data^36^. To experimentally test the hypothesis that disruption of *IDO1* impacts both tryptophan and kynurenine pool sizes, we used CRISPR-Cas9 mediated knockdown with sgRNAs targeted against *IDO1* human colorectal carcinoma HCT116 cells. These experiments corroborated earlier data indicating that knockdown of *IDO1* depleted kynurenine pools (**Figure 2F** and **Figure 2G**). However, while the association between *IDO1* and kynurenine has been widely described in the literature^36^, our analysis indicated that *IDO1* is also expected to be pool-determining for tryptophan, an amino acid involved in numerous other reactions in the cell, most obviously the synthesis of proteins (**Figure 2H**). In another experimental validation, we observed that knockdown of *IDO1* was sufficient to increase tryptophan levels, indicating that the pool size of this highly connected proteinogenic amino acid could be perturbed in part through disruption of *IDO1* activity (**Figure 2I**). Second, in support of a proximal GMI between *GGT1-* GSSG (**Figure 2J**), we reanalyzed existing metabolomic data from a functional knockdown of *GGT1* vs. control in human embryonic kidney HEK293T cells^42,43^. This data confirmed that knockdown of *GGT1* was associated with an increase in GSSG levels with respect to mock control (p-value = 2.90×10^−2^), suggesting that *GGT1* is a pool-determining consumer of GSSG (**Figure 2K**). Taken together, these data demonstrate that lineage-transcending GMIs discovered through pathway-based analysis of the cAMP represents examples of genes exerting strong control over metabolite pool sizes.

### Metabolic and transcriptomic changes are not coregulated at a pathway level

Despite the interesting findings related to proximal GMIs, the vast majority of GMIs in our datasets did not represent proximal pathway reactions. In total, only 2.5% (565/22,619) of GMIs were proximal, whereas 97.5% of statistically significant GMIs represented distant, non-proximal interactions beyond obvious enzyme-substrate metabolic relationships (**Figure 2B**). One possible implication of such non-local covariation is that genes and metabolites in the same metabolic pathway would show asynchronous and uncorrelated changes across different groups of samples, such as tumors and normal tissues. To investigate this hypothesis, we studied the consistency of transcriptional and metabolic differences in tumor versus adjacent normal tissue across cancer types. To this end, we performed differential analysis of metabolite and transcript levels between tumor and normal tissues in the 7 cAMP datasets where both tissues were available (**Table 1**) and aggregated the results into 85 KEGG metabolic pathways. Out of these, we considered 63 pathways with at least one metabolite or gene measured in at least 5 of the 7 cAMP datasets (**Table S2**).

For each KEGG pathway, we evaluated (using a differential abundance/”DA” score, see **Methods**) whether metabolites and transcripts showed synchronous accumulation or depletion patterns in tumors relative to normal tissues. Nevertheless, pathways were biased towards asynchronous changes (276/441, 63%), where increases in metabolite levels coincided with decreases in transcript levels, and vice versa (**Figure 3A**). Only one pathway (Histidine metabolism) demonstrated fully synchronous changes in all datasets, whereas a few others demonstrated uniformly asynchronous changes (*e*.*g*., Primary Bile Acid Biosynthesis). We also assessed whether there was a correlation between the extent of metabolomic versus transcriptomic disruption regardless of the direction (using a differential fraction/”DF” score, see **Methods**). A minority of pathways (9/63) demonstrated significant associations (nominal p-value < 0.05) between RNA and metabolite DF scores (**Figure 3B**, see **Figure 3C** as an example). Interestingly, these 9 pathways belonged to just two KEGG pathway classes (**Table S3**): Amino acid metabolism and Carbohydrate metabolism. Enrichment analysis indicated that the class of carbohydrate metabolism pathway (out of 8 total) was significantly over-represented relative to the others (Fisher’s exact test p-value: 2.36×10^−2^). Thus, the majority of pathways showed no evidence of a correlation between metabolomic and transcriptomic disruption, emphasizing the implications of predominantly distally-acting GMIs and prompting the question of which biological phenomena produce these distal GMIs.

**Figure 3.**
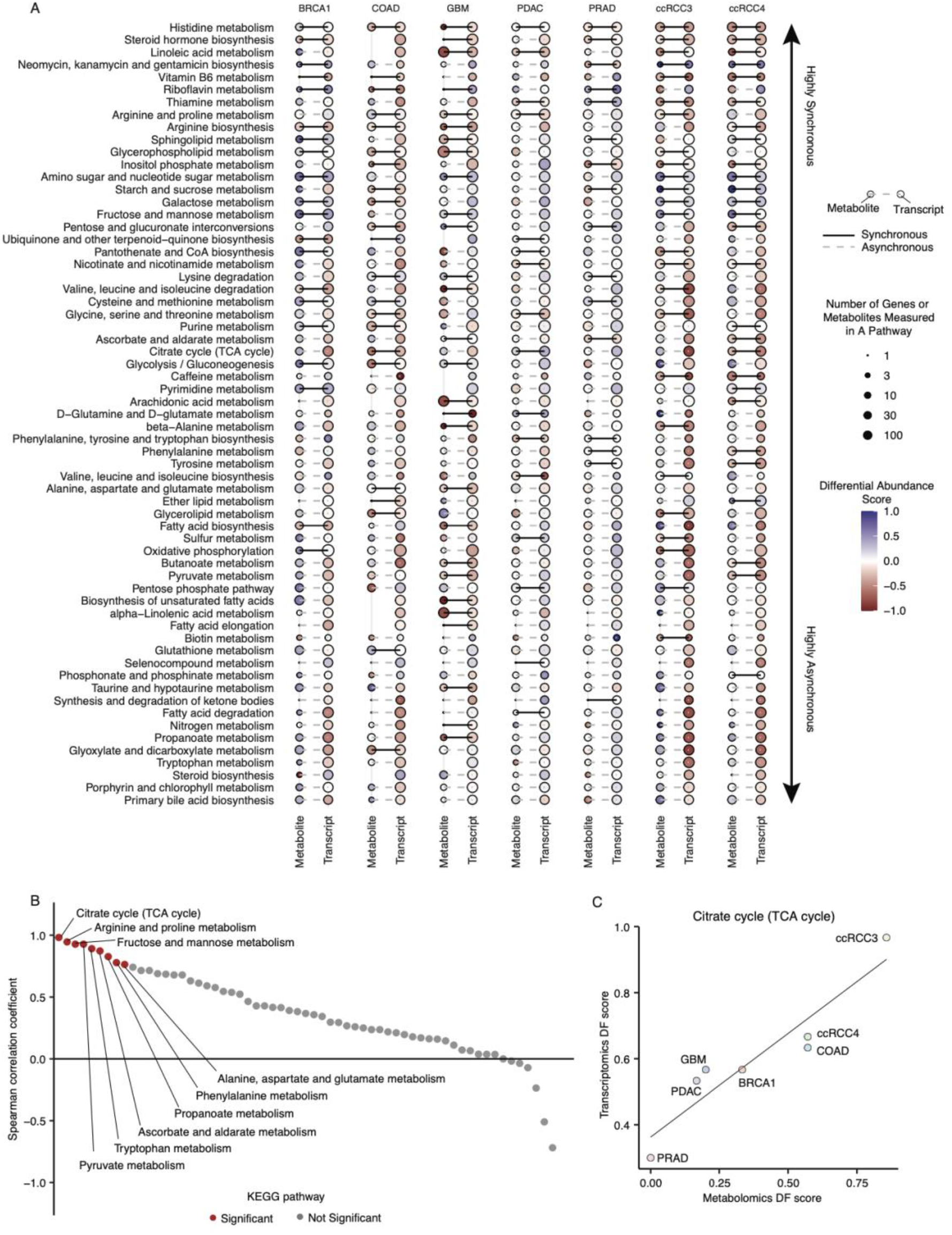
Tumor vs. normal changes in metabolite and transcript abundance are predominantly asynchronous. (A) Heatmap of metabolite and transcript differential abundance (“DA”) scores across datasets and pathways, capturing the tendency for metabolites and genes to accumulate or deplete in tumors relative to normal tissues. The size of the dots indicates the number of molecules measured in that pathway while the color represents the DA score. (B) Spearman correlation coefficients of the metabolite and transcript differential fraction (“DF”) scores in KEGG pathways. Red dots indicate nominal significance (p-value<0.05). A minority of pathways showed significant association between metabolomic and transcriptomic disruption across cAMP datasets. (C) Example of Spearman correlation calculation: Distribution of the metabolite (x-axis) and transcript (y-axis) DF scores across datasets for the Citrate cycle (TCA cycle) pathway.

### Hub metabolites are enriched for associations with immune genes

Having established that the majority of GMIs do not represent biochemically proximal interactions, we adopted a broader approach to identify the driving factors of metabolite-transcript correlations. First, we investigated the distribution of GMIs across metabolites and genes, observing that GMIs were strongly concentrated in a small number of metabolites (**Figure 4A**). The top three metabolites, quinolinate, nicotinamide mononuclueotide (NMN), and 5’-methylthioadenosine (MTA) alone contributed to 17% (3,823/22,619) of all GMIs in our analysis, and the top ten covered 35% of the GMIs (8,048/22,619), far higher than the fraction covered by the top 10 genes (**Figure 4B**). That is, a small number of metabolites participated in an exceptionally high number of GMIs, acting as “hubs” for strong co-variation with gene expression. Interestingly, hub metabolites concentrated in certain metabolic pathways: Among the top 10 most correlated metabolites, we found several constituents of the NAD+ biosynthesis pathway (quinolinate, NMN, NAD+) and nucleotide metabolism (thymine, uracil, adenine).

**Figure 4.**
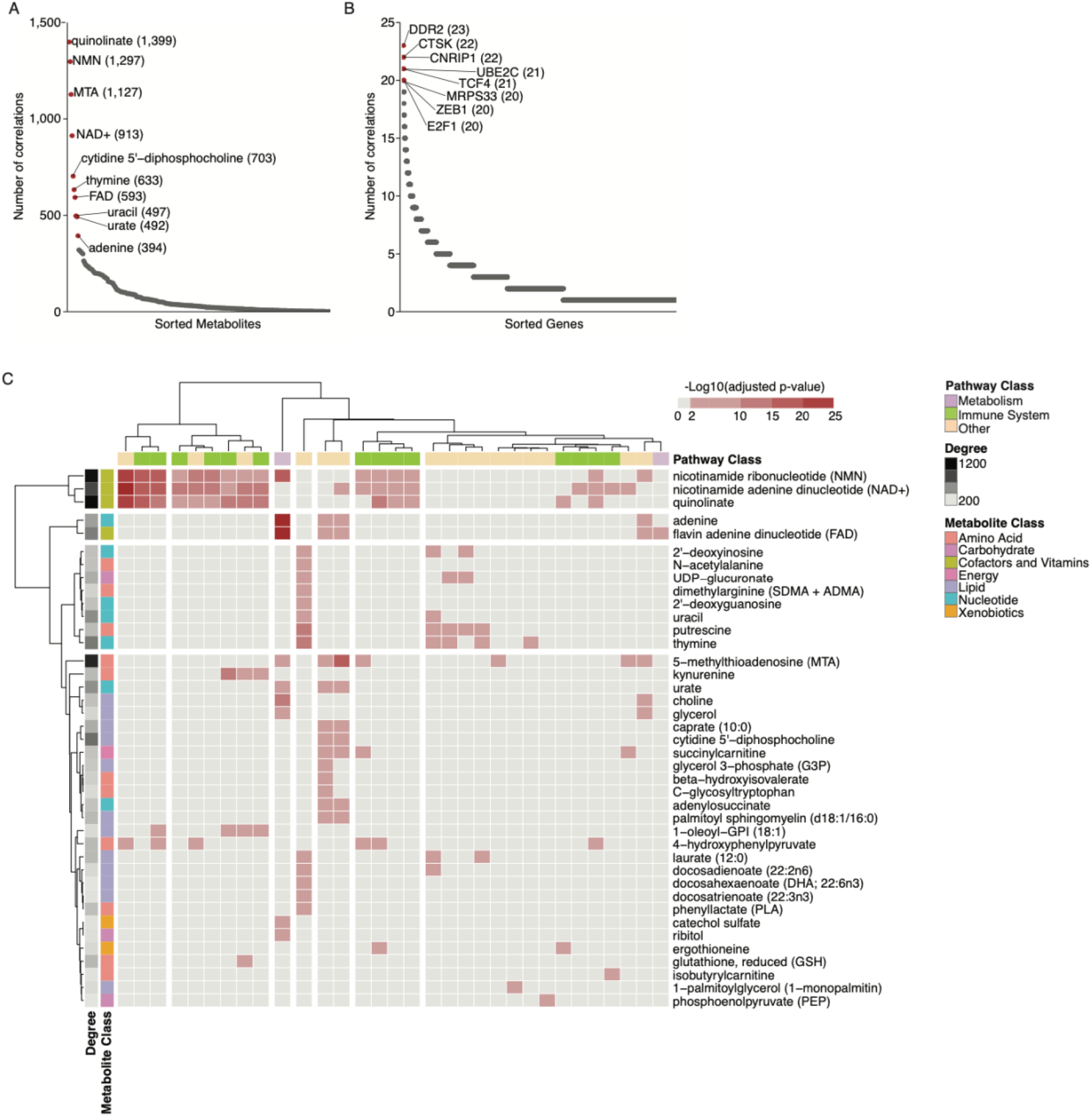
GMIs are concentrated in hub metabolites associated with immune-related genes. (A) Distribution of significant GMIs across metabolites. The top 10 metabolites with the highest number of significant associations are labeled. (B) Distribution of significant GMIs across genes. The top 10 genes with the highest number of significant associations are labeled. (C) KEGG-based gene pathway enrichment analysis results for metabolites with at least one GMI and at least one significantly enriched pathway. The heatmap color represents the strength of the enrichment as the negative log_10_(p-value) of the pathway enrichment test. Cells colored in shades of red indicate pathways that were significant after multiple testing correction (FDR<0.01), and gray cells were insignificant associations. Individual pathways were classified based on the type of process they describe: Metabolism (lilac), immune (green), or other (cream).

To determine whether the genes correlated with a particular metabolite were enriched for specific cellular functions, we performed an unsupervised pathway enrichment analysis. For each metabolite with at least one GMI, we investigated whether specific pathways and processes were over-represented in the transcripts correlated with that metabolite. Overall, the analysis used 146 KEGG pathways (**Table S4**), covering the 85 metabolic processes in **Figure 3** as well as cellular processes (e.g., cell growth and death), signaling pathways, genetic processing pathways (e.g., transcription and translation), and organismal systems pathways (e.g., immune, endocrine and sensory system). 32 unique pathways were over-represented across 40 metabolites (adjusted p-value < 0.01, **Figure 4C**). Interestingly, only two of those 32 pathways represented metabolic processes: oxidative phosphorylation, for which the top 3 most associated metabolites were adenine, NMN, and FAD, and the TCA cycle, also associated with FAD. The remaining top-ranked pathways were exclusively non-metabolic. Notably, three metabolite hubs in particular, quinolinate, NMN, and NAD+, showed broad enrichment for immune-related cellular processes, including chemokine signaling, as well as B-cell and T-cell receptor signaling pathways. This indicates that these metabolites, which are involved in the NAD+ biosynthetic pathway, are correlated with the expression of a wide array of genes associated with the immune response.

### NAD+-related metabolites are associated with immune cell infiltration across cancers

Human tumor tissues are heterogeneous compositions of various cell populations, including tumor cells, immune cells, and stromal cells. Bulk and single cell profiling technologies have revealed that a large subset of genes are exclusively expressed in immune cells or non-immune cell subpopulations *(e*.*g*., tumor cells)^44,45^. We reason that the correlation between metabolites in the NAD+ biosynthesis pathway and immune-related genes (**Figure 4C**) could therefore arise if NAD+-related metabolites were at a characteristically higher or lower abundance in immune cells relative to non-immune cells. One implication of this hypothesis is that, while each cancer type might demonstrate its own unique metabolomic changes associated with immune infiltration, NAD+-related metabolites should be at elevated concentrations in highly immune-infiltrated tumors relative to lowly-infiltrated tumors across many different cancer types.

To determine if NAD+-related metabolites were associated with immune infiltration across tumors, we used single-sample gene set enrichment (ssGSEA)^46^ to compute a previously validated 141-gene RNA signature of overall immune cell infiltration (“ImmuneScore”) directly from bulk RNA sequencing data^47^, and identified the individual metabolites correlated with this immune phenotype. Concordance between metabolite levels and the ImmuneScore signature was assessed across all samples in each cAMP dataset (**Figure 5A**). In general, covariation between specific metabolite pools and ImmuneScore expression was cancer-type specific. For example, of the 12 metabolites significantly associated with ImmuneScore in intrahepatic cholangiocarcinoma (ICC) and hepatocellular carcinoma (HCC) (representing the top 2 cancer types with the highest number of metabolites significantly associated to ImmuneScore), only 3 metabolites were consistently associated with ImmuneScore in both datasets (**Figure 5B, Table S5**). Notably, we did not observe a significant correlation (Spearman’s rank correlation p-value = 0.69) between the percentage of metabolites significantly associated with ImmuneScore and expression of the ImmuneScore signature itself (shown in **Figure 5A**, central panel).

**Figure 5.**
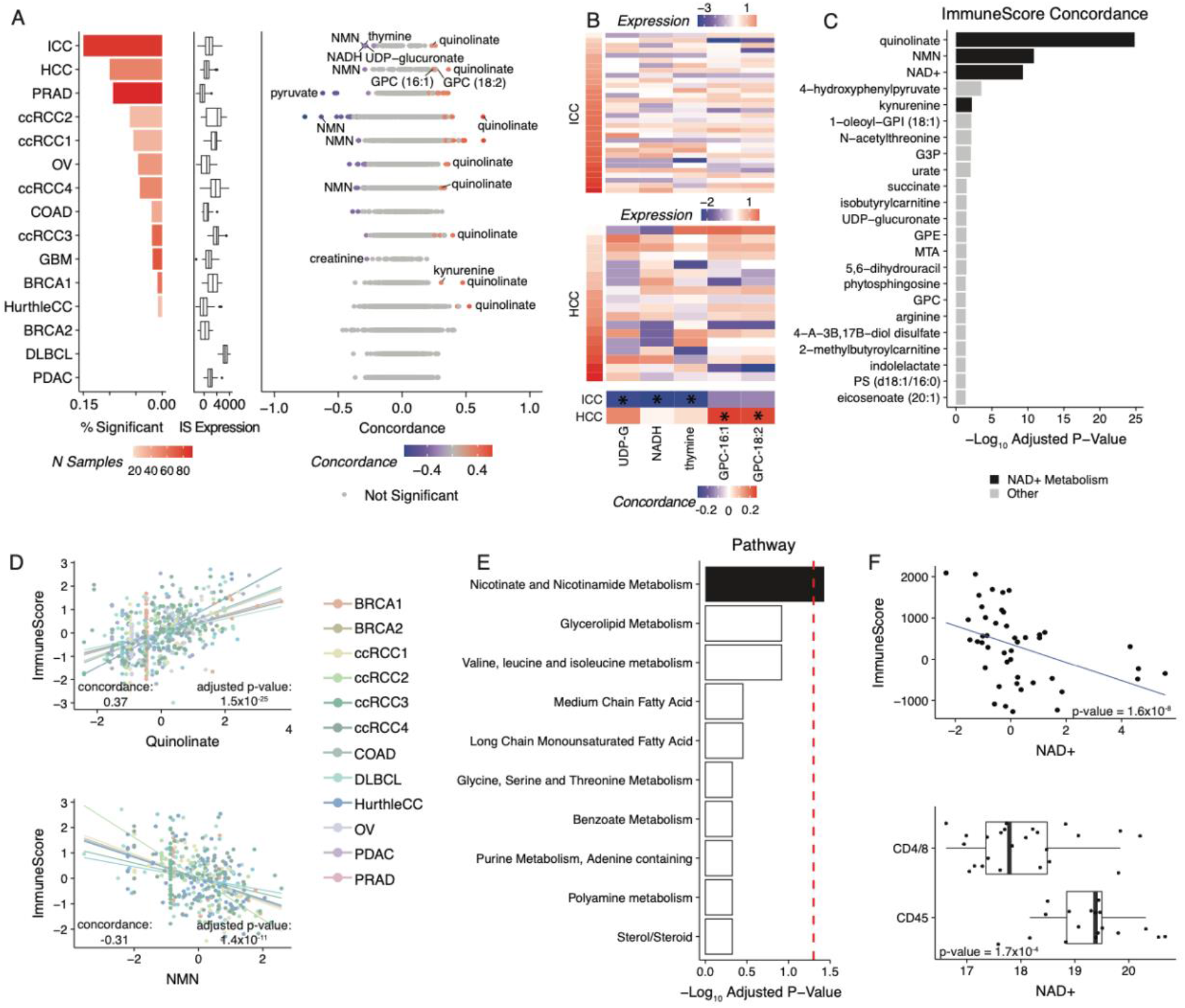
The abundance of NAD+-related metabolites is shaped by immune infiltration in the tumor microenvironment. (A) Left: Association of metabolite abundance with immune infiltration varies significantly across cancer types. The barplot indicates the fraction of metabolites significantly associated with the ImmuneScore signature in each dataset. Bar colors code for the sample size of the dataset. Middle: Boxplots indicate the expression range of the ImmuneScore signature within each dataset. Right: Plot of scaled concordance calculated between metabolites and ImmuneScore. Red dots indicate metabolites with a positive association with ImmuneScore, blue dots metabolites with negative associations, and gray dots indicate metabolites that were not statistically significant. (B) Heatmaps of five metabolites, three of which are highly correlated to ImmuneScore in the ICC dataset and two which are highly correlated with ImmuneScore in the HCC dataset. Samples are sorted by increasing ImmuneScore. UDP-G = UDP-glucuronate, GPC-16:1 = 1-palmitoleoyl-GPC (16:1), GPC-18:2 = 1-linoleoyl-GPC (18:2). (C) Bar plot indicating the strength of association between metabolites and ImmuneScore from concordance meta-analysis. Bar length represents the -log_10_ adjusted concordance meta-analysis p-value. Metabolites related to NAD+ metabolism are shown in black. (D) Scatterplots of the abundance of two NAD+-related metabolites, quinolinate (top) and NMN (bottom), versus ImmuneScore expression across all datasets. Metabolite abundances were scaled within each dataset. (E) Bar plot of FDR-adjusted p-values from a one-sided Wilcoxon test comparing the absolute concordance values of metabolites to ImmuneScore in a pathway compared to all other pathways. (F) Metabolomic measurements of purified populations of CD45-tumor cells and CD45+ T cells isolated from ovarian cancer tumors. NAD+ was negatively correlated to the ImmuneScore signature in our ovarian cancer dataset. NAD+ was similarly lower in abundance in CD4/8+ cells than CD45-(non-immune) cells in the dataset of purified cell populations.

While metabolite concordance with ImmuneScore was generally heterogeneous across cAMP datasets, we observed that several NAD+-related metabolites, including quinolinate and NMN, were recurrently associated with high or low levels of immune infiltration in numerous disease contexts (**Figure 5A**). To systematically identify such lineage-agnostic metabolomic correlates of immune infiltration, we again applied concordance meta-analysis, identifying 23 metabolites significantly associated with ImmuneScore across datasets (adjusted p-value< 0.05, **Figure 5C**), with quinolinate and NMN being the strongest hits (**Figure 5D**). Interestingly, 4/14 significantly associated metabolites (quinolinate, NAD+, NMN, and kynurenine) were members of the NAD+ biosynthesis pathway. Consistent with this, we identified NAD+ metabolism as the sole pathway whose metabolites demonstrated significantly stronger association with immune infiltration than all other pathways (adjusted p-value: 0.04, **Figure 5E**).

The above analysis suggested that NAD+-related metabolites displayed a differential abundance profile in immune cells relative to non-immune cells, and that this effect produced a consequent accumulation of NAD+-related metabolites in immune-infiltrated tumors. Some support for this hypothesis can be found in previously published immunohistochemical data indicating that the abundance of quinolinate increases dramatically in diverse immune cell populations in response to Toll-like receptor 4 (TLR4) ligands such as lipopolysacharide (LPS)^48^. To provide more evidence for this hypothesis, we compared our findings to a recently published study of metabolomic profiles of purified CD45-tumor cells and CD45+ (CD8+ and CD4+) T cells from ovarian cancer tumors^48^. In our data, NAD+ was negatively correlated with ImmuneScore in ovarian cancer, suggesting that it was at lower abundance in immune cells relative to non-immune cells (**Figure 5F**). Consistent with this, NAD+ was at significantly lower abundance in CD45+ T cells than CD45-tumor cells in the dataset of purified cell populations (NAD+ Log_2_FC = −1.22, p value = 1.7×10^−4^) (**Figure 5F**). Together, these analyses suggest that the pool sizes of NAD+-related metabolites are at characteristically different abundance in immune cells relative to other cell types, and that this effect ultimately drives the association of quinolinate and other NAD+-related metabolites with immune infiltration in bulk tumor data.

### Kynurenine and histamine accumulate in the presence of specific immune cell populations

Whereas in the previous analysis we investigated the association of metabolite levels with overall immune infiltration, we next turned to investigating the association of metabolite levels and specific immune cell populations *(e*.*g*., T cells, macrophages, and numerous other cell types), each with unique transcriptional phenotypes and immunologic functions^49–51^. To investigate how these diverse immune cell populations contributed to the observed GMIs, we estimated the abundance of 23 immune cell types from bulk transcriptomics profiles^51^ using ssGSEA as shown previously^46^, and computed their association to metabolite levels across cancers. We again focused on lineage-agnostic relationships by performing a concordance meta-analysis to calculate associations between metabolite levels and immune cell signatures across tumors from all cancer types. In total, 7.3% of all metabolite-signature pairs (466 out of 6,348 pairs) demonstrated statistically significant associations (adjusted p-value < 0.05, **Figure 6A**). Among these, quinolinate was positively associated with almost all immune cell populations (17/23), consistent with prior immunohistochemical data and suggesting that it accumulates in a variety of immune cell types in a cancer-type-agnostic manner^52^.

**Figure 6.**
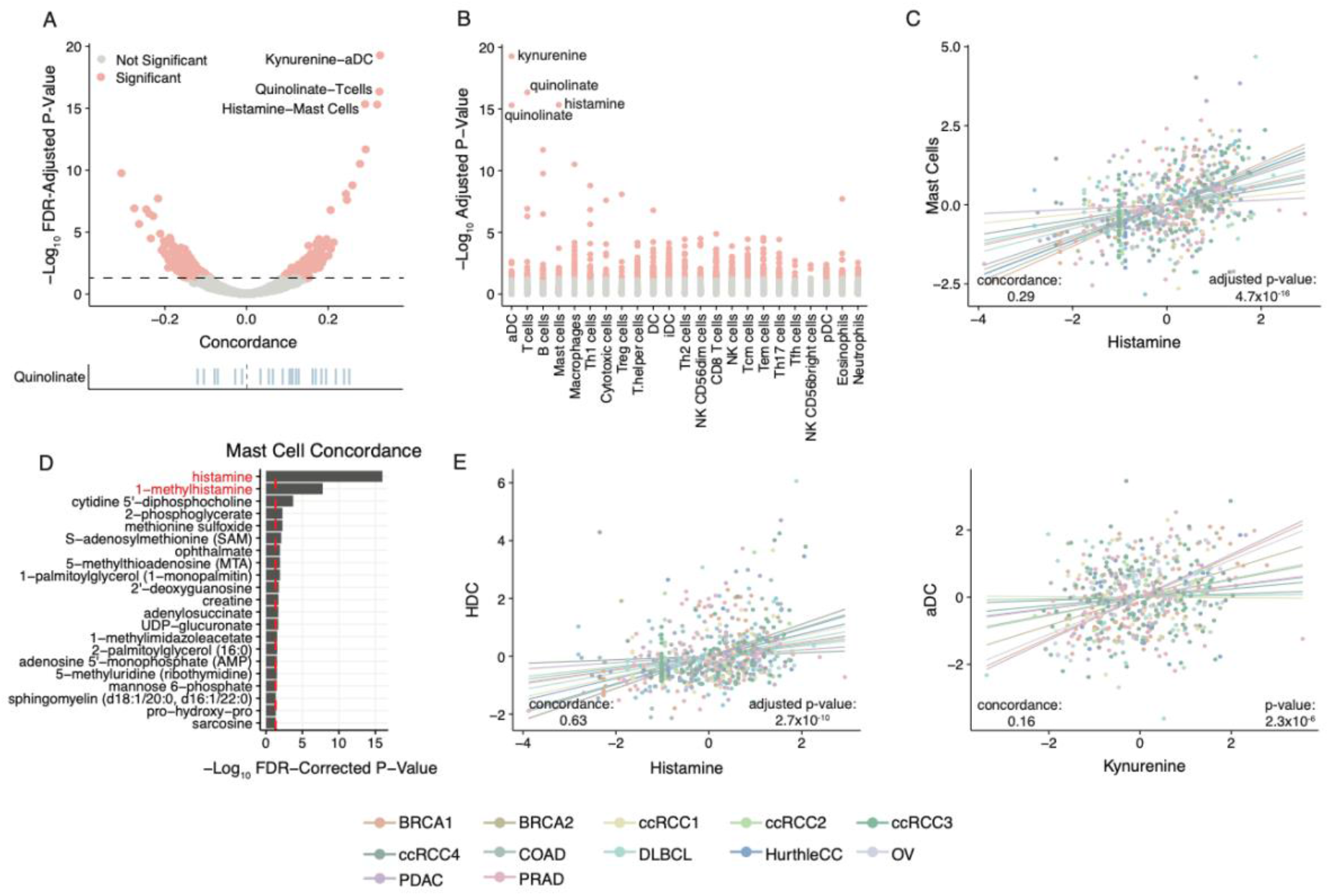
A subset of metabolites associates with specific immune cell lineages. (A) Volcano plot of cell type signature-metabolite interactions. The rug plot at the bottom highlights the numerous statistically significant associations with quinolinate. (B) Manhattan plot of adjusted concordance p-values between metabolites and cell types. (C) Histamine associates with the abundance of mast cells across most datasets of the cAMP. (D) Bar plot of adjusted concordance p-values of metabolites correlated with the mast cell gene signature. Red dotted line indicates the significance cutoff of 0.05. (E) *HDC* expression strongly associates with mast cell abundance across the cAMP. (F) Kynurenine abundance associates with an activated dendritic cell signature across the cAMP.

Aside from quinolinate, the two strongest associations between immune cell type signatures and metabolite levels were two comparatively rare cell populations, namely mast cells-histamine and activated dendritic cells (aDCs)-kynurenine (**Figure 6B**). Mast cells are a myeloid cell population which, when stimulated, mediate the inflammatory process by synthesizing histamine from histidine using the enzyme *HDC*. We found both histamine and its related metabolite 1--methylhistamine were the two metabolites most significantly associated with the presence of mast cells (**Figure 6C** and **Figure 6D**), and that the association between histamine and mast cell abundance was likely driven by diverse cancer types in the cAMP (**Figure S2**). Histamine levels themselves were strongly associated with the expression of *HDC* across the cAMP (**Figure 6E**). Interestingly, the median variation of histamine across the tumor datasets in the cAMP was ∼740-fold, implying that fluctuations in the abundance of an otherwise rare cell population were sufficient to produce large-magnitude changes in the abundance of a histamine in the bulk tumor.

In contrast to mast cells and their physiological role in producing histamine, dendritic cells are not known to be dedicated sources of kynurenine in the microenvironment, although single cell data indicates that *IDO1* expression is strongly elevated in dendritic cells relative to other cell types (**Figure S3**). While the aDC signature contains *IDO1* (and *IDO1* participates in a strong GMI with kynurenine, **Figure 2**), an aDC signature without *IDO1* preserved strong positive concordance with kynurenine (concordance = 0.59, p-value = 2.3×10^−6^), confirming that this association was not solely dependent on *IDO1* itself (**Figure 6F**). *IDO1* and kynurenine corresponded to the strongest GMI in the cAMP, raising two opposing hypotheses on the mechanism underlying the primary source of kynurenine in the tumor microenvironment: either bulk tumor expression of *IDO1* is primarily determined by high dendritic-cell-specific expression, or alternatively, it is driven by a comparatively low expression in far more abundant non-dendritic-cell populations. Resolving this ambiguity would require single cell measurements of metabolite levels which have recently become technically feasible^53^. Together, the data above suggests that the presence of individual and comparatively rare cell populations is associated with shifts in the abundance of immunomodulatory metabolites in the tumor microenvironment.

## Discussion

Metabolism is jointly controlled by genetically-encoded enzymes and small molecule metabolites. To study the interactions between genes and metabolites at scale, we assembled and harmonized a database of metabolomic and transcriptomic data from ∼1,000 tumor and normal samples which we refer to as the Cancer Atlas of Metabolic Profiles (cAMP). Although large-scale multimodal measurements of metabolism have previously been produced in bacteria^54^ and yeast^55^, a comparable resource for human cancers had been missing. The cAMP is among the first such resources and represents a significant public database for studying the metabolism of complex human tissues and cancers. Our analysis of the cAMP demonstrates that large-scale studies of multimodal metabolic data can reveal fundamental principles of metabolic regulation both at the scale of individual metabolic reactions (**Figure 2**) as well as complex human tissues (**Figure 5** and **Figure 6**).

Our detailed statistical analysis of the cAMP revealed a multitude of gene-metabolite associations that transcended the tissue of origin. We focused on two specific types of gene-metabolite co-regulation. The first, induced by functional proximity, identified metabolic genes whose activation or inhibition is likely to have a significant effect on the direct substrate or product of the respective reaction (**Figure 2**). While metabolite pools are likely controlled by a large number of genes and other factors, our data-driven analysis identified a small subset of metabolites whose pool size was strongly associated to a single, proximal gene across tissue lineages. The GMIs identified in **Figure 2B** offer a data-informed, rational approach for modulating the pool sizes of their metabolite constituents. While each of the highlighted metabolites in **Figure 2B** may participate in numerous metabolic reactions, data from the cAMP specifically nominates single genes (*e*.*g*., *GGT1* for GSSG, and *IDO1* for tryptophan and kynurenine) as those targets whose perturbation is most likely to disrupt the corresponding pool size.

The second broad form of gene-metabolite coregulation, likely induced by cell-type-specific physiology, corresponded to metabolites whose abundance was associated with the presence of specific immune cells in the microenvironment. This class of GMIs was characterized by a small number of metabolites (enriched for NAD+-related molecules) that were significantly correlated with a large number of genes, and in particular immune-related genes. This apparent association between the abundance of specific immune cells in a tissue specimen and the levels of numerous NAD+-associated metabolites suggests that immune cells have evolved mechanisms to maintain the concentrations of these metabolites at characteristically different levels relative to other cell types. Because NAD+ is both the central mediator of redox poise in the cell and a cofactor for numerous metabolic and non-metabolic reactions, understanding the mechanisms by which immune cells achieve differential abundance of NAD+-related metabolites, and the selective pressure to do so, can provide insights into the metabolic phenotypes underlying both cancer and other diseases involving dysfunctional immune responses^56^. Given the growing use of spatial metabolomics technologies that can measure the metabolomic profiles of individual cells in heterogenous tissue slices^57–60^, the full extent of cell-type-specific metabolomic alterations seems now in experimental reach.

Discoveries about the function of cancer genes have emerged from a combination of untargeted, population-scale genomic surveys^61^ and mechanistic experiments in specific disease and genetic backgrounds^62^. Combining these approaches has proven transformative for the discovery of recurrent and large-effect-size alterations and prompted their characterization in model systems of human disease. In contrast, the field of cancer metabolism has primarily been driven by bottom-up experiments with modest support from large-scale (largely genetic or transcriptomic) datasets^63–65^. The cAMP is a counterpoint to these efforts: By assembling and harmonizing in one database metabolomic and transcriptomic data from diverse diseases, the cAMP represents a unique opportunity for *de novo* discovery of translationally relevant metabolic phenotypes in cancer. Expanding the scope of the cAMP to include other forms of data, including but not limited to genomic sequencing, epigenetic profiling, and proteomic measurements, holds the potential to reveal entirely new and highly recurrent metabolic phenomena in cancer.

## Methods

### Collection of Cancer Atlas of Metabolic Profiles (cAMP)

We combined 12 published datasets with 3 additional in-house datasets that profiled metabolite and gene expression from the same samples to create a comprehensive collection of 988 samples (764 tumor sample and 264 adjacent normal samples) across 11 different cancer types, covering 15 datasets, which we called the Cancer Atlas of Metabolic Profiles (cAMP). Details and references associated with these studies are provided in **Table 1** and **Figure 1A**. Data are available for download under DOI: 10.5281/zenodo.7150252.

### Gene Expression Data Processing Pipeline

Six of the cAMP datasets included gene expression data captured by the Affymetrix platform (GSE28735, GSE37751, GSE26193, GSE62452, GSE76297, and Cornell_PROSTATE). For these datasets, CEL files were downloaded from GEO or from their source within our respective institutions. Then, we applied the Robust Multichip Average (RMA) algorithm for background subtraction, quantile normalization, and summarization (via median-polish) by using the “rma” function implemented in the R oligo package (version 1.40.2)^66^. Each dataset’s rma-normalized expression matrix was then used for downstream analysis. In the COAD dataset, gene expression data was captured by an Agilent custom array, and we downloaded the gene expression matrix deposited at the GEO repository (GSE89076) for further downstream analysis.

For cAMP datasets with RNA sequencing data, RNA-seq reads were aligned against human genome assembly hg19 by STAR 2-pass alignment^67^ (version 2.5.3a). QC metrics, such as general sequencing statistics, gene feature, and body coverage, were then calculated based on the alignment result through RSeQC^68^ (version 2.6.4). RNA-seq gene level count values were computed by using the R package GenomicAlignments^69^ (version 1.14.2) over aligned reads with UCSC KnownGene^70,71^ with hg19 as the base gene model. The Union counting mode was used and only mapped paired reads after alignment quality filtering were considered. Finally, gene level FPKM (Fragments Per Kilobase Million) and raw read count values were computed by the R package DESeq2^72^ (version 1.18.1).

Across the 15 cAMP datasets, 16,082 out of ∼40,000 distinct transcripts were profiled in all cohorts and used for analysis (**Figure S4A**).

### Metabolomics Data Preprocessing

The metabolomics data for 3/15 datasets (GBM, LiCa1 and LiCa2) were provided preprocessed and were therefore used in their original form. OV data was already normalized and was only log_2_-transformed before analysis. For the other 11 metabolomics datasets in our study, we obtained the raw metabolomics data from the data owners. In this case, the processing pipeline was standardized across datasets and included batch correction via median scaling if multiple batches of data were present (only necessary for PRAD dataset as it was the only dataset produced in distinct batches) and probabilistic quotient normalization^73^ using either only normal samples if available, or all tumor samples if no normal sample was included, and only metabolites with less than 20% values missing to create the reference sample. After normalization, metabolite abundances were log_2_-transformed. Metabolites with more than 80% values missing were excluded from the analysis. For the remaining metabolites, missing values were imputed using the minimum value recorded. Data preprocessing was performed using the R package maplet. Metabolite names and annotations were manually harmonized for consistency by matching HMDB^74^, KEGG^75^ and Metabolon platform-specific peak IDs across datasets. The overlap of metabolites across datasets was heterogeneous (**Figure S4B**): Out of 2,411 unique molecules quantified across all datasets, fewer than 500 were measured in more than 5 datasets, and only three metabolites (gluconate, glucose, and glucose-3-phosphate) were quantified in all 15 datasets in tumor. This high variability can be attributed to a variety of technical and biological factors, including metabolite ionizability on the mass spectrometer and potential specificity of certain metabolites to distinct tissues.

### Concordance meta-analysis

To identify gene-metabolite pairs that were consistently associated across tumor types and cohorts, we used a stratified, weighted concordance model. Concordance is a non-parametric measure of correlation, similar to Kendall’s Tau, that relies on the concept of concordant pairs^37^.

Briefly, consider a pair of observations *i* and *j*, for which both metabolite *m* and gene *g* have been measured. The pair is defined to be *concordant* if *sign(m*_*i*_ − *m*_*j*_*) = sign(g*_*i*_ − *g*_*j*_), that is, if they have the same order in both samples, and *discordant* if they have opposite signs. Once pairwise concordance has been estimated for all pairs, the overall concordance *c* is calculated as

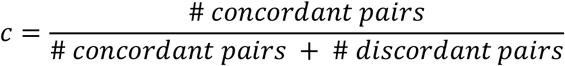

To account for the multiple cohorts in our study, we performed a *stratified* concordance analysis, where pairwise comparisons are only calculated within datasets, but not across. A global concordance value is then estimated by counting the overall number of concordant pairs according to the formula above. Moreover, given the vast heterogeneity in sample sizes across studies, we downweighed each observation by the number of samples in the corresponding study (i.e., 1*/nsamples(dataset*_*i*_)), so that each dataset would contribute equally to the overall concordance. The concordance calculation was performed using the concordance function from the survival R package (v3.2-3)^76^.

To make this quantity more intuitive, we further scaled the concordance range to values between −1 and 1 through *c*_*scaled*_ = 2*c* −1, *which is analogous to Somers’ D*^77^. Furthermore, in the absence of ties, this value is also identical to Kendall’s rank correlation coefficient *tau*^77^. In the figures presented in this paper, a scaled concordance of 0 indicates absence of association, a value greater than 0 indicates positive association, while a value less than 0 indicates negative association.

A z-score was computed as concordance minus 0.5 and divided by the square root of the variance, and the resulting value was used to derive a two-tailed p-value^78,79^. P-values were corrected for multiple testing using the Benjamini-Hochberg method to control the false discovery rate^80^. Pairs with adjusted p-values below 0.01 were considered significant.

### Generation of *IDO1* Knockdown HCT116 Cell Line

sgRNAs (oligonucleotide sequences are indicated in **Table S6**) targeting *IDO1* as well as non-targeting control were cloned into lentiCRISPRv2 puro plasmid (Addgene, #98290). Lentiviral packaging vectors psPAX2 (Addgene, #12260), pMD2.G (Addgene, #12259), along with sgRNA expressing vector were transfected into HEK293T cells using PEI (polyethyleneimine) transfection reagent. 72 hours after transfection, supernatant containing lentivirus was harvested and filtered through a 0.45 μm pore size Whatman filter (Fisher Scientific) to remove cell debris. Target cells (HCT116, human colon cancer cell line, ATCC) were transduced with lentivirus using 8 μg/ml polybrene (Sigma). 72 hours after lentivirus transduction, 3 μg/ml puromycin (ThermoFisher) was added to cell culture media to select for virus-infected cells. 2 weeks after puromycin selection, target gene knock-out was confirmed by western blot. For induction of *IDO1* gene expression, cells were treated with 100 ng/ml IFN-γ for 48 hours.

### Metabolomics Profiling on the *IDO1* Knockdown HCT116 Cell Line

Cells were plated in 6-well tissue culture plates at 100K cells/well density. 72 hours after cell seeding, metabolites were extracted and analyzed by liquid chromatography-mass spectrometry (LC-MS). For metabolite extraction, culture media was aspirated, and cells were washed once with ice-cold phosphate buffered saline (PBS). After PBS washing, 1 ml of ice-cold extraction solvent (methanol:water = 80:20) was added. After overnight incubation at −80°C, cells and extraction solvent were collected into 1.5ml Eppendorf tube using cell scraper. Samples were centrifuged at 20,000*g* for 20 min at 4°C. Supernatant (900 μl) was collected and dried in a vacuum evaporator (Genevac EZ-2 Elite).

For LC-MS, dried extracts were resuspended in 30 μl of 97:3 water:methanol containing 10 mM tributylamine and 15 mM acetic acid. Samples were vortexed, incubated on ice for 20 min, and clarified by centrifugation at 20,000*g* for 20 min at 4°C. LC-MS analysis used a Zorbax RRHD Extend-C18 column (150 mm x 2.1 mm, 1.8 μm particle size, Agilent Technologies). Solvent A was 10 mM tributylamine, 15 mM acetic acid in 97:3 water:methanol, and solvent B was 10 mM tributylamine and 15 mM acetic acid in 3:97 water: methanol, prepared according to the manufacturer’s instructions (MassHunter Metabolomics dMRM Database and Method, Agilent Technologies). LC separation was coupled to a 6470 triple quadrupole mass spectrometer (Agilent Technologies) which was operated in dynamic MRM scan type and negative ionization mode.

### Tumor vs Normal Pathway Analysis

There were 7 cAMP datasets which had both tumor and normal samples available (BRCA1, COAD, GBM, PRAD, PDAC, ccRCC3 and ccRCC4). We applied differential gene expression tests between tumor and normal samples in each dataset using limma-voom (limma package version: 3.5.2). Genes with an FDR-adjusted p-value < 0.1 were considered significantly differentially expressed. Similarly, we also conducted differential metabolite abundance testing between tumor and normal samples using Wilcoxon rank sum tests. Metabolites with an FDR-adjusted p-value below 0.1 were considered significantly differentially abundant.

For each KEGG pathway, we calculated the differential fraction (“DF”) score and the differential abundance (DA) score for genes and metabolites separately:

DF score = (number of significantly up-regulated constituents + number of significantly down-regulated constituents) / number of measured constituents in a pathway

DA score = (number of significantly up-regulated constituents - number of significantly down-regulated constituents) / number of measured constituents in a pathway

Constituents are either genes or metabolites in the above formula.

Conceptually, the DF score captures the overall disruption of the constituents of a pathway, whereas the DA score captures the tendency for pathway constituents to increase or decrease in abundance relative to a reference (in this case, normal tissue) state.

### Gene-Metabolite Distance

We used a genome-scale human metabolic model (“Human 1” model^41^) covering the metabolic reaction information of transporters, enzymes and metabolites to calculate the distance between gene and metabolite pairs. We defined a “proximal” interaction by a gene/metabolite pair participating in the same reaction (shortest path distance of 1) or participating in subsequent reactions (shortest path distance of 2) (**Figure S5**).

### Gene Set Enrichment Analysis

For the analysis in **Figure 4**, we ran a pathway enrichment analysis among all significant genes for each metabolite with at least one significant gene association. The enrichment test was performed using classical hypergeometric testing^81^. We tested the enrichment of 146 KEGG pathways. P-values were adjusted using the Benjamini-Hochberg method for controlling the false discovery rate^80^. Adjusted p-values < 0.01 were considered significant.

Results were then aggregated into a metabolite x pathway matrix and visualized as a heatmap, where metabolites and pathways were clustered based on –log_10_(p-value). Clustering was performed using the pheatmap function^82^ (pheatmap package v 1.0.12) with Ward linkage and Euclidean distance.

### Bulk Gene Expression Deconvolution Analysis

To dissect the role of the immune compartment in the tumor microenvironment, we calculated the ImmuneScore through the estimate R package^83^. To calculate cell-type-specific infiltration patterns in Figure 6, we employed the ssGSEA^46^ for bulk gene expression deconvolution analysis. Signature gene lists of immune cell types as well as immune features were obtained from Bindea et al.^51^. Briefly, ssGSEA takes the sample FPKM expression values as the input and computes an enrichment score for a given gene list as compared to all the other genes in the sample transcriptome.

## Acknowledgments

We thank each of the individual laboratories which contributed their data to the cAMP. The Reznik Lab acknowledges Payam Gammage for useful feedback and discussions.

## Supplementary Table Legends

**Table S1**.

Table of statistically significant GMIs and associated statistics.

**Table S2**

Summary of differential abundance results for pathways considered in tumor vs. normal analysis.

**Table S3**

Association of DF gene/metabolite scores across pathways in tumor vs. Normal analysis.

**Table S4**

Pathways considered for the analysis in Figure 4.

**Table S5**

Comparison of metabolite correlated to ImmuneScore in the HCC and ICC datasets.

**Table S6**

Guide RNA sequences used for *IDO1* knockout experiment.

## References

1. Pavlova, N. N. & Thompson, C. B. The emerging hallmarks of cancer metabolism. Cell Metab. 23, 27–47 (2016).

2. Zhang, J., Nuebel, E., Daley, G. Q., Koehler, C. M. & Teitell, M. A. Metabolic regulation in pluripotent stem cells during reprogramming and self-renewal. Cell Stem Cell 11, 589– 595 (2012).

3. Intlekofer, A. M. & Finley, L. W. S. Metabolic signatures of cancer cells and stem cells. Nat. Metab. 1, 177–188 (2019).

4. Fanciulli, M. et al. Energy Metabolism of Human LoVo Colon Carcinoma Cells: Correlation to Drug Resistance and Influence of Lonidamine1 | Clinical Cancer Research | American Association for Cancer Research. Clinical Cancer Research (2000).

5. Zhao, Y., Butler, E. B. & Tan, M. Targeting cellular metabolism to improve cancer therapeutics. Cell Death Dis. 4, e532 (2013).

6. Zhou, Y. et al. Intracellular ATP levels are a pivotal determinant of chemoresistance in colon cancer cells. Cancer Res. 72, 304–314 (2012).

7. O’Sullivan, D., Sanin, D. E., Pearce, E. J. & Pearce, E. L. Metabolic interventions in the immune response to cancer. Nat. Rev. Immunol. 19, 324–335 (2019).

8. Hurley, H. J. et al. Frontline Science: AMPK regulates metabolic reprogramming necessary for interferon production in human plasmacytoid dendritic cells. J. Leukoc. Biol. 109, 299–308 (2021).

9. Guerra, L., Bonetti, L. & Brenner, D. Metabolic modulation of immunity: A new concept in cancer immunotherapy. Cell Rep. 32, 107848 (2020).

10. Domblides, C., Lartigue, L. & Faustin, B. Control of the antitumor immune response by cancer metabolism. Cells 8, (2019).

11. Ganeshan, K. & Chawla, A. Metabolic regulation of immune responses. Annu. Rev. Immunol. 32, 609–634 (2014).

12. Anders, S. et al. Count-based differential expression analysis of RNA sequencing data using R and Bioconductor. Nat. Protoc. 8, 1765–1786 (2013).

13. Penney, K. L. et al. Metabolomics of prostate cancer gleason score in tumor tissue and serum. Mol. Cancer Res. 19, 475–484 (2021).

14. Gentric, G. et al. PML-Regulated Mitochondrial Metabolism Enhances Chemosensitivity in Human Ovarian Cancers. Cell Metab. 29, 156-173.e10 (2019).

15. Chaisaingmongkol, J. et al. Common molecular subtypes among asian hepatocellular carcinoma and cholangiocarcinoma. Cancer Cell 32, 57-70.e3 (2017).

16. Wang, L.-B. et al. Proteogenomic and metabolomic characterization of human glioblastoma. Cancer Cell 39, 509-528.e20 (2021).

17. Calvo-Vidal, M. N. et al. Oncogenic HSP90 Facilitates Metabolic Alterations in Aggressive B-cell Lymphomas. Cancer Res. 81, 5202–5216 (2021).

18. Satoh, K. et al. Global metabolic reprogramming of colorectal cancer occurs at adenoma stage and is induced by MYC. Proc Natl Acad Sci USA 114, E7697–E7706 (2017).

19. Terunuma, A. et al. MYC-driven accumulation of 2-hydroxyglutarate is associated with breast cancer prognosis. J. Clin. Invest. 124, 398–412 (2014).

20. Tang, X. et al. A joint analysis of metabolomics and genetics of breast cancer. Breast Cancer Res. 16, 415 (2014).

21. Zhang, G. et al. Integration of metabolomics and transcriptomics revealed a fatty acid network exerting growth inhibitory effects in human pancreatic cancer. Clin. Cancer Res. 19, 4983–4993 (2013).

22. Hakimi, A. A. et al. An integrated metabolic atlas of clear cell renal cell carcinoma. Cancer Cell 29, 104–116 (2016).

23. Warburg, O. On the Origin of Cancer Cells. Science 123, 309–314 (1956).

24. Garcia-Bermudez, J. et al. Aspartate is a limiting metabolite for cancer cell proliferation under hypoxia and in tumours. Nat. Cell Biol. 20, 775–781 (2018).

25. Krall, A. S. et al. Asparagine couples mitochondrial respiration to ATF4 activity and tumor growth. Cell Metab. 33, 1013-1026.e6 (2021).

26. Martínez-Reyes, I. et al. Mitochondrial ubiquinol oxidation is necessary for tumour growth. Nature 585, 288–292 (2020).

27. Sullivan, L. B. et al. Aspartate is an endogenous metabolic limitation for tumour growth. Nat. Cell Biol. 20, 782–788 (2018).

28. Labuschagne, C. F., van den Broek, N. J. F., Mackay, G. M., Vousden, K. H. & Maddocks, O. D. K. Serine, but not glycine, supports one-carbon metabolism and proliferation of cancer cells. Cell Rep. 7, 1248–1258 (2014).

29. Yang, M. & Vousden, K. H. Serine and one-carbon metabolism in cancer. Nat. Rev. Cancer 16, 650–662 (2016).

30. Puccetti, P. et al. Accumulation of an endogenous tryptophan-derived metabolite in colorectal and breast cancers. PLoS ONE 10, e0122046 (2015).

31. Yaku, K., Okabe, K., Hikosaka, K. & Nakagawa, T. NAD metabolism in cancer therapeutics. Front. Oncol. 8, 622 (2018).

32. Chiarugi, A., Dölle, C., Felici, R. & Ziegler, M. The NAD metabolome--a key determinant of cancer cell biology. Nat. Rev. Cancer 12, 741–752 (2012).

33. Cancer Genome Atlas Research Network. Comprehensive genomic characterization defines human glioblastoma genes and core pathways. Nature 455, 1061–1068 (2008).

34. Finlay, D. & Cantrell, D. A. Metabolism, migration and memory in cytotoxic T cells. Nat. Rev. Immunol. 11, 109–117 (2011).

35. Auslander, N. et al. A joint analysis of transcriptomic and metabolomic data uncovers enhanced enzyme-metabolite coupling in breast cancer. Sci. Rep. 6, 29662 (2016).

36. Li, H. et al. The landscape of cancer cell line metabolism. Nat. Med. 25, 850–860 (2019).

37. Therneau, T. M. & Grambsch, P. M. Modeling survival data: extending the cox model. (Springer New York, 2000). doi:10.1007/978-1-4757-3294-8.

38. Badawy, A. A.-B. Kynurenine pathway of tryptophan metabolism: regulatory and functional aspects. Int. J. Tryptophan Res. 10, 1178646917691938 (2017).

39. Chini, C. C. S. et al. CD38 ecto-enzyme in immune cells is induced during aging and regulates NAD+ and NMN levels. Nat. Metab. 2, 1284–1304 (2020).

40. Hitchings, G. H. & Falco, E. A. The identification of guanine in extracts of girella nigricans: the specificity of guanase. Proc Natl Acad Sci USA 30, 294–297 (1944).

41. Robinson, J. L. et al. An atlas of human metabolism. Sci. Signal. 13, (2020).

42. Priolo, C. et al. Impairment of gamma-glutamyl transferase 1 activity in the metabolic pathogenesis of chromophobe renal cell carcinoma. Proc Natl Acad Sci USA 115, E6274–E6282 (2018).

43. Long, Y. et al. Liver-Specific Overexpression of Gamma-Glutamyltransferase Ameliorates Insulin Sensitivity of Male C57BL/6 Mice. J. Diabetes Res. 2017, 2654520 (2017).

44. Reinfeld, B. I. et al. Cell-programmed nutrient partitioning in the tumour microenvironment. Nature 593, 282–288 (2021).

45. Zappasodi, R. et al. CTLA-4 blockade drives loss of Treg stability in glycolysis-low tumours. Nature 591, 652–658 (2021).

46. Barbie, D. A. et al. Systematic RNA interference reveals that oncogenic KRAS-driven cancers require TBK1. Nature 462, 108–112 (2009).

47. Yoshihara, K. et al. Inferring tumour purity and stromal and immune cell admixture from expression data. Nat. Commun. 4, 2612 (2013).

48. Kilgour, M. K. et al. 1-Methylnicotinamide is an immune regulatory metabolite in human ovarian cancer. Sci. Adv. 7, (2021).

49. Qian, B.-Z. & Pollard, J. W. Macrophage diversity enhances tumor progression and metastasis. Cell 141, 39–51 (2010).

50. Biswas, S. K. & Mantovani, A. Macrophage plasticity and interaction with lymphocyte subsets: cancer as a paradigm. Nat. Immunol. 11, 889–896 (2010).

51. Bindea, G. et al. Spatiotemporal dynamics of intratumoral immune cells reveal the immune landscape in human cancer. Immunity 39, 782–795 (2013).

52. Moffett, J. R. et al. Quinolinate as a marker for kynurenine metabolite formation and the unresolved question of NAD+ synthesis during inflammation and infection. Front. Immunol. 11, 31 (2020).

53. Rappez, L. et al. SpaceM reveals metabolic states of single cells. Nat. Methods 18, 799– 805 (2021).

54. Fuhrer, T., Zampieri, M., Sévin, D. C., Sauer, U. & Zamboni, N. Genomewide landscape of gene-metabolome associations in Escherichia coli. Mol. Syst. Biol. 13, 907 (2017).

55. Mülleder, M. et al. Functional metabolomics describes the yeast biosynthetic regulome. Cell 167, 553-565.e12 (2016).

56. Navas, L. E. & Carnero, A. NAD+ metabolism, stemness, the immune response, and cancer. Signal Transduct. Target. Ther. 6, 2 (2021).

57. Taylor, M. J., Lukowski, J. K. & Anderton, C. R. Spatially resolved mass spectrometry at the single cell: recent innovations in proteomics and metabolomics. J. Am. Soc. Mass Spectrom. 32, 872–894 (2021).

58. Abbas, I. et al. Kidney lipidomics by mass spectrometry imaging: A focus on the glomerulus. Int. J. Mol. Sci. 20, (2019).

59. Sun, N. et al. Mass Spectrometry Imaging Establishes 2 Distinct Metabolic Phenotypes of Aldosterone-Producing Cell Clusters in Primary Aldosteronism. Hypertension 75, 634– 644 (2020).

60. Basu, S. S. et al. Rapid MALDI mass spectrometry imaging for surgical pathology. NPJ Precis. Oncol. 3, 17 (2019).

61. Chang, M. T. et al. Accelerating discovery of functional mutant alleles in cancer. Cancer Discov. 8, 174–183 (2018).

62. Findlay, G. M. et al. Accurate classification of BRCA1 variants with saturation genome editing. Nature 562, 217–222 (2018).

63. Nilsson, R. et al. Metabolic enzyme expression highlights a key role for MTHFD2 and the mitochondrial folate pathway in cancer. Nat. Commun. 5, 3128 (2014).

64. Gatto, F., Nookaew, I. & Nielsen, J. Chromosome 3p loss of heterozygosity is associated with a unique metabolic network in clear cell renal carcinoma. Proc Natl Acad Sci USA 111, E866–75 (2014).

65. Hu, J. et al. Heterogeneity of tumor-induced gene expression changes in the human metabolic network. Nat. Biotechnol. 31, 522–529 (2013).

66. Carvalho, B. S. & Irizarry, R. A. A framework for oligonucleotide microarray preprocessing. Bioinformatics 26, 2363–2367 (2010).

67. Dobin, A. et al. STAR: ultrafast universal RNA-seq aligner. Bioinformatics 29, 15–21 (2013).

68. Wang, L., Wang, S. & Li, W. RSeQC: quality control of RNA-seq experiments. Bioinformatics 28, 2184–2185 (2012).

69. Lawrence, M. et al. Software for computing and annotating genomic ranges. PLoS Comput. Biol. 9, e1003118 (2013).

70. Harrow, J. et al. GENCODE: the reference human genome annotation for The ENCODE Project. Genome Res. 22, 1760–1774 (2012).

71. Harrow, J. et al. GENCODE: producing a reference annotation for ENCODE. Genome Biol. 7 Suppl 1, S4.1-9 (2006).

72. Love, M. I., Huber, W. & Anders, S. Moderated estimation of fold change and dispersion for RNA-seq data with DESeq2. Genome Biol. 15, 550 (2014).

73. Dieterle, F., Ross, A., Schlotterbeck, G. & Senn, H. Probabilistic quotient normalization as robust method to account for dilution of complex biological mixtures. Application in 1H NMR metabonomics. Anal. Chem. 78, 4281–4290 (2006).

74. Wishart, D. S. et al. HMDB: the human metabolome database. Nucleic Acids Res. 35, D521–6 (2007).

75. Kanehisa, M. & Goto, S. KEGG: Kyoto encyclopedia of genes and genomes. Nucleic Acids Res. 28, 27–30 (2000).

76. Therneau, T. M. A Package for Survival Analysis in R. https://CRAN.R-project.org/package=survival (2022).

77. Newson, R. Parameters behind “Nonparametric” Statistics: Kendall’s tau, Somers’ D and Median Differences. The Stata Journal 2, 45–64 (2002).

78. Therneau, T. M. & Watson, D. A. The concordance statistic and the Cox model. Technical Report #85 1–18 (2017).

79. Newson, R. Confidence intervals for rank statistics: somers’ D and extensions. The Stata Journal 6, 309–334 (2006).

80. Benjamini, Y. & Hochberg, Y. Controlling the false discovery rate: a practical and powerful approach to multiple testing. Journal of the Royal Statistical Society: Series B (Methodological) 57, 289–300 (1995).

81. Reimand, J. et al. Pathway enrichment analysis and visualization of omics data using g:Profiler, GSEA, Cytoscape and EnrichmentMap. Nat. Protoc. 14, 482–517 (2019).

82. Kolde, R. CRAN - Package pheatmap. https://cran.r-project.org/web/packages/pheatmap/index.html (2019).

83. Yoshihara, K., Kim, H. & Verhaak, R. G. estimate: Estimate of Stromal and Immune Cells in Malignant Tumor Tissues from Expression Data. https://r-forge.r-project.org/projects/estimate/ (2016).

84. Han, Y. et al. TISCH2: expanded datasets and new tools for single-cell transcriptome analyses of the tumor microenvironment. Nucleic Acids Res. (2022) doi:10.1093/nar/gkac959.

85. Sun, D. et al. TISCH: a comprehensive web resource enabling interactive single-cell transcriptome visualization of tumor microenvironment. Nucleic Acids Res. 49, D1420–D1430 (2021).

86. Ganly, I. et al. Mitonuclear genotype remodels the metabolic and microenvironmental landscape of Hürthle cell carcinoma. Sci. Adv. 8, eabn9699 (2022).

