## Supplementary Figures for "A Multimodal Atlas of Tumor Metabolism Reveals the Architecture of Gene-Metabolite Co-regulation"

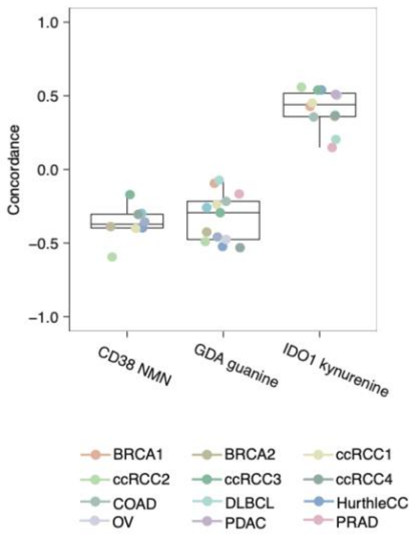


**Figure S1.** Box plots of concordance for gene-metabolite pair of CD38-MNM, GDA-guanine and IDO1-kynurenine across individual datasets.

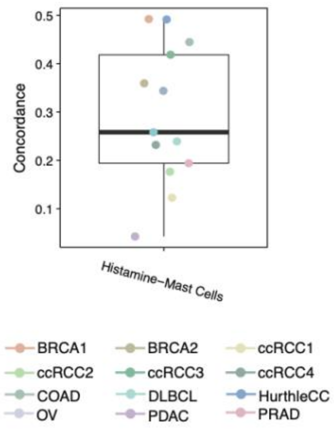


**Figure S2.** Box plots of concordance of histamine – mast cells across individual datasets.

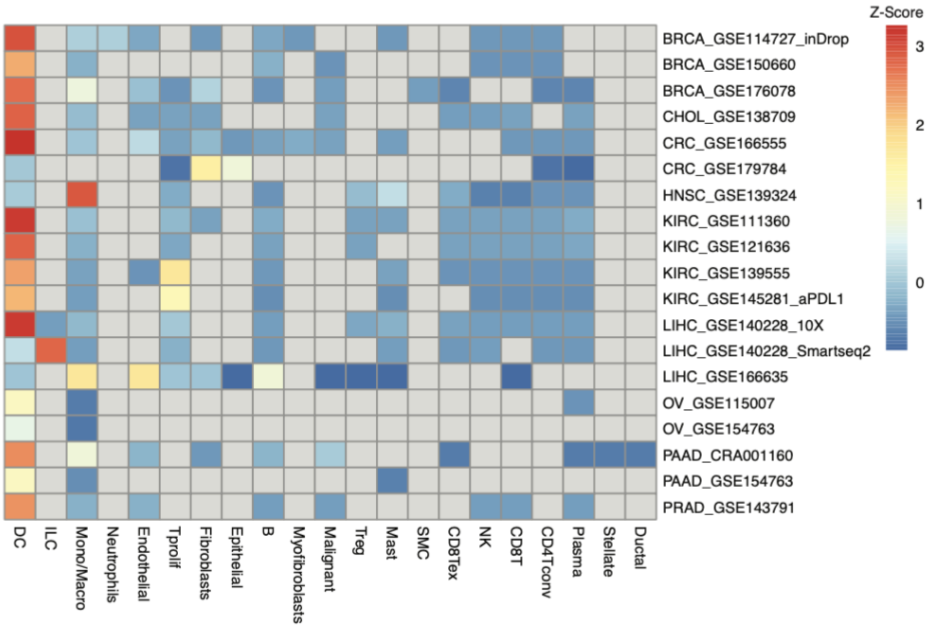


**Figure S3.** Heatmap of single cell gene expression data for *IDO1* across datasets from the TISCH2 database[84,85]. Expression is standardized to z-scores across cell types in a dataset. Considering only datasets in which dendritic cells are identified, dendritic cells exhibit the highest expression of *IDO1*.

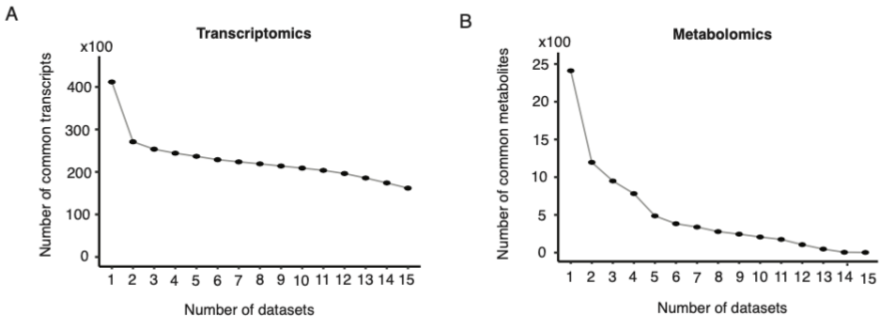


**Figure S4.** (A) Number of common transcripts across datasets in tumor tissue. (B) Number of common metabolites across datasets in tumor tissue.

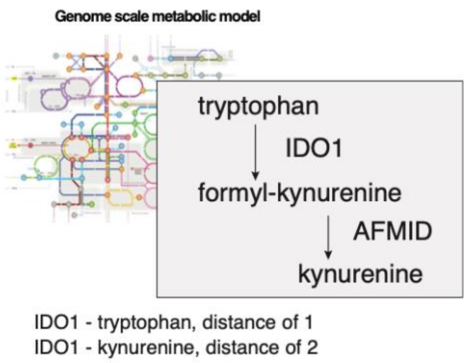


**Figure S5.** Definition of “proximal” interactions in the Human 1 metabolic network model, illustrated by the example of IDO1 distances to tryptophan and kynurenine.
